# Lifespan Trajectories of Alpha Rhythm: Dynamic Shifts in Neural Excitation-Inhibition Balance

**DOI:** 10.1101/2025.05.14.653960

**Authors:** Chuanliang Han, Vincent C.K. Cheung, Rosa H.M. Chan

## Abstract

Alpha rhythm (8-13 Hz), a key neural oscillation in the brain, plays a significant role in cognitive functions and reflects the brain’s excitatory-inhibitory (E-I) balance. This study investigates the dynamics of alpha rhythm across the lifespan, focusing on how E-I balance modulates alpha power and peak frequency, and exploring the distinct age-related and sex-specific patterns of alpha activity. Using a computational E-I model, we simulated the impact of different neuronal connections and E-I ratios on alpha rhythm characteristics. The results suggest that self-regulation primarily affects alpha power, while interaction between excitatory and inhibitory neurons influences both alpha frequency and power. We applied this model to real EEG data from 3265 participants across a wide age range, revealing that alpha power and peak frequency exhibit an inverted U-shape across the lifespan, peaking in early adulthood and declining in old age. Significant sex differences in alpha activity were observed primarily during puberty and later in life. Decomposition of the alpha band into periodic and aperiodic components showed that periodic activity follows the inverted U-shape, while aperiodic activity declines exponentially with age. Our findings indicate that alpha rhythm is governed by complex E-I dynamics, with distinct contributions from periodic and non-periodic components, and highlight the role of alpha rhythm in age-related cognitive changes and sex differences in brain function.

## Introduction

Alpha rhythm, typically ranging from 8 to 13 Hz, is among the most prominent rhythms observed in the human brain^1–5^ and have been linked to various cognitive functions^6–10^, including attention^11–16^, memory^17–25^ and learning^26–29^. Fluctuations in alpha-band activity reflects changes in the brain’s excitability and functional states. Recent research suggests that the modulation of alpha rhythms is closely linked to the delicate balance between neural excitation and inhibition (E-I balance)^30–32^, which is fundamental for proper cognitive functions^33–40^. Despite the importance of alpha rhythms in brain function, the underlying mechanisms driving their modulation are not fully understood.

During brain development and aging, the E-I balance undergoes significant shifts^41,42^, which would influence the performance in behavioral level, and then may affect characteristics of alpha rhythms. In behavioral studies, classic psychological research has found that human performance on cognitive tasks follows a U-shape (or inverted U-shape) pattern throughout life^43–46^. Human performance starts to improve gradually during childhood, peaks in early adulthood, and declines with aging. This trajectory of behavioral development is thought to be related to inhibition functions, as human inhibitory abilities increase with age, peaking in adulthood and weakening in old age. The classic theory of alpha rhythms in the brain is input suppression^47,48^, meaning they are greatly enhanced in the closed-eye state and weakened in the open-eye state, with similar results observed during spatial attention tasks^49–51^. Alpha rhythm is associated with inhibitory functions as well, but how its oscillatory characteristics evolve across the human lifespan remains unclear, as well as that for excitatory functions. If alpha rhythms are indeed related to behavioral performance, they should also follow a U-shape curve, becoming stronger and then weaker with age. However, this has yet to be verified.

Alpha rhythm naturally follows specific patterns with age^52–54^, but sex is also an important variable that needs to be considered^55–59^. Sex differences in brain function and structure are well-documented across species^60^, with animal studies uncovering specific neural mechanisms^61–63^. However, how these differences manifest in EEG oscillatory features and the underlying neural mechanisms remains unclear. Previous studies investigating sex differences in EEG rhythms have yielded inconsistent results^59^, especially across alpha band ^64–69^, likely due to limited sample sizes, age confounds, or unclear behavioral correlates. Hence, how sex differences in alpha rhythm evolves across the lifespan still remains unclear.

This study aims to explore the developmental trajectory of oscillatory properties of alpha rhythm (alpha power and peak frequency) across the lifespan and its neural mechanism behind based on computational models and real large-scale EEG datasets (n=3265). By integrating findings from multiple EEG datasets, and computational models, we seek to elucidate how the evolution of alpha rhythm develops across different stages of life. Additionally, we examined the role of sex differences in shaping these rhythms and investigate how spatial dynamics in some specific brain regions, such as the occipital and parietal areas, relate to these changes. In the end, we provided a more comprehensive understanding of the neural mechanisms that govern alpha rhythms and their role in cognitive development and aging.

## Materials and Methods

### Public EEG dataset - Healthy Brain Network (HBN) Dataset^70^

The EEG data was collected from 2951 participants (age from 5 to 22, Male, Female) during resting state (eyes-open (EO) and eyes-closed (EC)). The public dataset could be download in the following link: https://openneuro.org/ datasets/ds004186/versions/2.0.0. The public dataset has been preprocessed; the details could also be seen in the link. The EEG was recorded with 128 electrodes and the total time length of closed-eye state is 40 seconds. The study was approved by the Chesapeake Institutional Review Board.

### Public EEG dataset - Max Planck Institute (MPI) Leipzig Mind-Brain-Body Dataset^71^

The EEG data was collected from 203 participants (age from 20 to 80, 129 Male, 74 Female) during resting state (eyes-open (EO) and eyes-closed (EC)). The public dataset could be download in the following link: https://fcon_1000.projects.nitr c.org/indi/retro/MPI_LEMON.html. The public dataset has been preprocessed; the details could also be seen in the above link. The EEG was recorded with 61 electrodes and the total time length of open- and closed-eye state is 8 minutes. The study was carried out in accordance with the Declaration of Helsinki and the study protocol was approved by the ethics committee at the medical faculty of the University of Leipzig (reference number 154/13-ff).

### Public EEG dataset - SRM Dataset^72^

The EEG dataset comprises resting-state recordings from 111 participants (age from 17 to 71, 42 Male, 69 Female) during eyes-closed (EC) conditions. Data were collected using a 64-electrode montage for a total duration of 4 minutes per participant. Detailed preprocessing steps were provided in the dataset documentation. The preprocessed dataset is publicly available at https://openneuro.org/datasets/ds003775/versions/1.2.1/download. The data collection was conducted in accordance with the Declaration of Helsinki, and informed consent was obtained from all participants. The procedures were approved by the Regional Ethics Committee of South-Eastern Norway (reference number: 2016/2003).

### Merging HBN, MPI and SRM dataset

MPI and SRM dataset are 64-electrode system, and HBN dataset is 128-electrode system. Hence, we combined them together to further analysis. By aligning two datasets in common electrodes, 58 electrodes are used in this study. It is noted that MPI dataset did not provide a specific age for each participant but instead gave an age range (such as 20-25, 25-30, 70-75). Therefore, in this study, we divided it into thirteen age range groups (5-6 years, 131M 65F; 6-7 years, 210M 104F; 7-8 years, 243M 118F; 8-9 years, 223M 144F; 9-10 years, 233M 118F; 10-11 years, 197M 98F; 11-12 years, 149M 85F; 12-14 years, 240M 118F; 14-16 years, 135M 100F; 16-18 years, 106M 67F; 18-30 years, 139M 91F; 30-55 years, 31M 41F; 55-80 years, 39M 40F) for comparison.

### Data Analysis

Data processing was performed in MATLAB (www.mathworks.com) with custom scripts. Data in three dataset was filtered between 1 and 30 Hz. The reference of all EEG dataset (MPI, HBN, SRM) was set to the averaged reference. We used spectrum analysis to quantify the neural oscillation strength in all electrodes in eyes-closed (EC) state. The power spectral density (PSD) for time series EEG data each electrode was computed using the multi-taper method with 5 tapers using the Chronux toolbox^73^, which is an open-source, data analysis toolbox available at http://chronux.org. Similar methods have been applied in various biomedical fields, such as experimental neuroscience^74^, neuropsychological disorders^75^, etc. Relative power was calculated by dividing the power at a specific frequency by the total power summation from 3 to 30Hz.

### Descriptive Model for Dissecting Aperiodic and Periodic Activity

The power spectrum was considered as the summation of two component: aperiodic and periodic components. The aperiodic component of the power spectrum was extracted using a 1/f like function (Equation 1), where a, b, c and d represent the model parameters to be estimated. The periodic component was defined as the residual obtained by subtracting the aperiodic component from the raw power spectrum. This method has been used in the characterization of oscillatory activities of different frequency bands^76–79^.

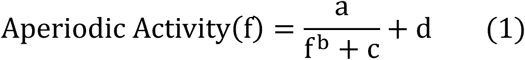

### Dynamical Model

We constructed a model unit, as previously described^80,81^. The model unit had two local components – local excitatory (E) and inhibitory (I) components^82–84^. The local components E and I can be thought of as a group of neurons recurrently connected (τ_E_=30ms, τ_#_=75ms). The dynamic interactions of E and I in local recurrent connection (RC) are described by Equations 2–4. The strengths of local interactions between E and I are denoted by *W*_$%_, where R denotes the receiver and S denotes the sender. The interaction type (excitatory or inhibitory) is denoted by the sign of *W*_$%_ (positive or negative).

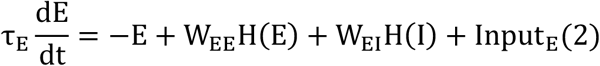

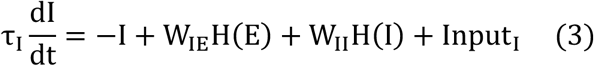

Where

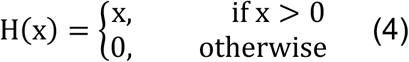

In the E-I unit, the local excitatory (E) component is connected to the local inhibitory (I) component with a coupling strength of W_IE_, and the connection strength within the E components is denoted by W_EE_. Similarly, the local inhibitory (I) component connects to the local excitatory (E) component with strength W_EI_, while the inhibitory connections within the I components are represented by W_II_. The time constants *τ*_&_ and *τ*_’_ in equations 2–3 correspond to the E and I components, respectively. The local field potential (LFP) is defined as the value of the E component at the central position (or in each local unit). The “spiking” thresholds for both the E and I components are set to 0 (as described by the function H(x) in Equation 4): only values of E and I that exceed this threshold (i.e., those greater than zero) will influence other neurons (i.e., H(E)). This can also be interpreted as the mean firing rate for a group of E or I neurons. Both the E and I components receive independent inputs from other brain regions or external sources, modeled as the mean of Gaussian white noise (std=1, Input_E_=80, Input_I_=40) ^85^, from which a random variable is drawn at each time step.

### Statistical Analysis

In merged HBN, MPI and SRM dataset, to screen out significant frequency bands and electrode positions for sex difference, the independent t-test was used to test difference between relative alpha power (Figure 3A) and alpha peak frequency (Figure 3B) of male and females in 13 age groups. Targeting on relative alpha power in electrodes in occipital region (O1, Oz, O2) that have large sex difference, two-way ANOVA (gender (M and F) and age (13 age groups) was conducted to relative power in closed-eye state respectively. Same method is also used in parietal region (P1, P2) for relative alpha power, and in parietal region (P1, P2) for alpha peak frequency.

## Results

The generation of alpha rhythm is essentially the result of interactions between different types of neuronal connections in neural circuits (Figure 1A). These interactions can be categorized into two types within a simple E-I model: self-regulation (connections within the same type of neurons, such as excitatory-to-excitatory W_EE_ or inhibitory-to-inhibitory W_II_) and interactive regulation (connections between different types of neurons, such as excitatory-to-inhibitory W_IE_ or inhibitory-to-excitatory W_EI_). The strength of the connections between different types of neurons determines the E-I balance of the circuit itself. When this balance is disrupted, the characteristics of the alpha rhythm in the brain also change.

**Figure 1.**
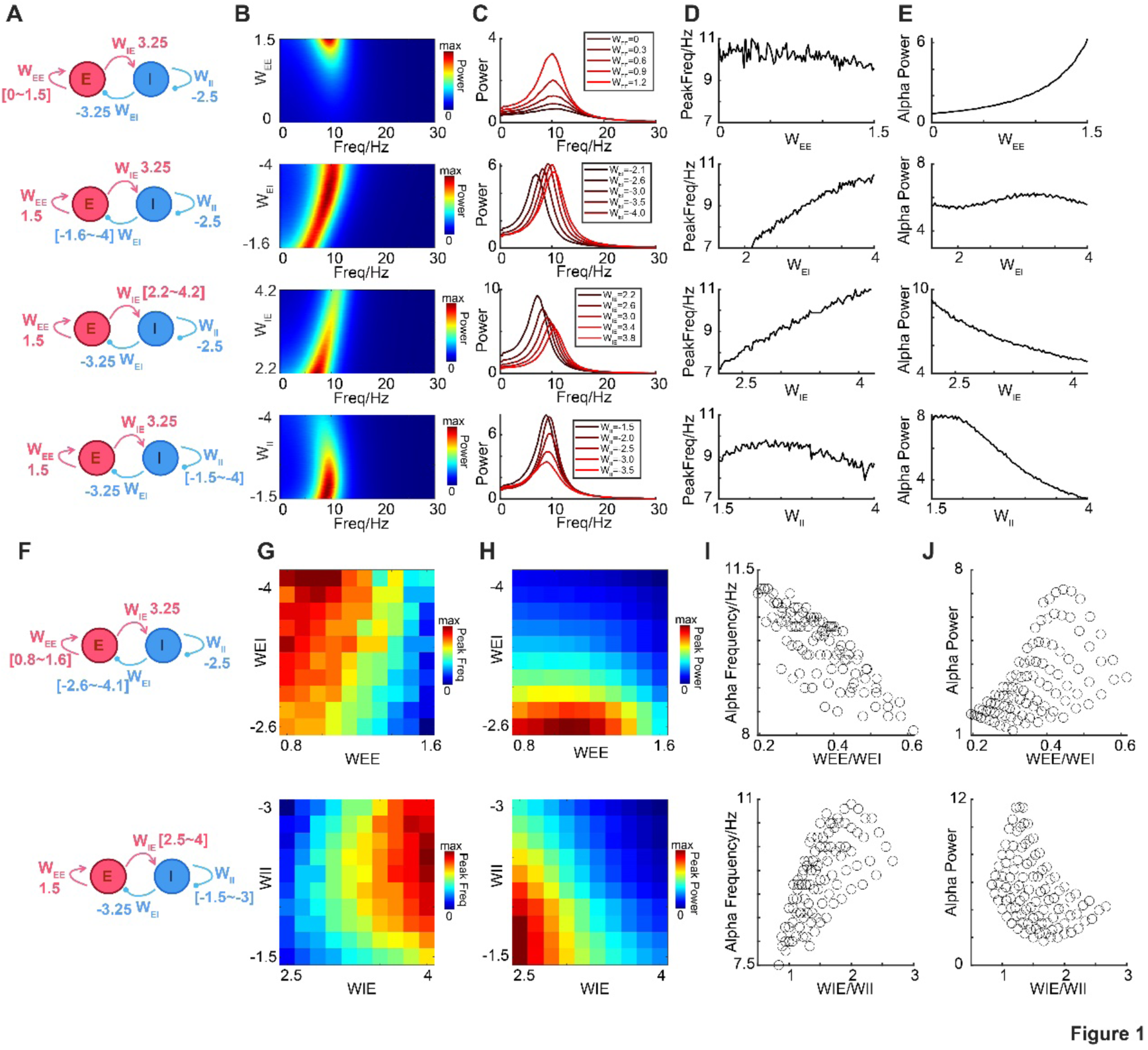
**A dynamic model illustrating the influence of E-I balance on alpha power and peak frequency.** A. Demo of E-I with different parameters. The range of parameters to be traversed is shown in square brackets. B. Modulation of power spectrum in alpha band by disturbing WEE, WEI, WIE and WII. Alpha power is shown in color. C. Modulation of power spectrum in alpha band by disturbing WEE, WEI, WIE and WII. D. Modulation of alpha peak frequency by disturbing WEE, WEI, WIE and WII. E. Modulation of alpha power by disturbing WEE, WEI, WIE and WII. F. Demo of E-I with different parameters. The range of parameters to be traversed is shown in square brackets. G. Modulation of alpha peak frequency by disturbing WEE/ WEI, and WIE/ WII. H. Modulation of alpha power by disturbing WEE/ WEI, and WIE/ WII. I. Scatter plot between alpha peak frequency and E-I ratios. J. Scatter plot between alpha power and E-I ratios.

### E-I balance shapes properties of alpha-band activity

To investigate the impact of different types of connections in neural circuits on alpha rhythm characteristics, we first employed a control variable method in this E-I model. Specifically, we sequentially perturbed four major parameters (sweeping one parameter while keeping others constant) to examine how the alpha-band activity in the spectrum is affected (Figure 1BC), especially for the peak frequency and power in alpha band (Figure 1EF). We found that self-regulation primarily affects the power of the spectrum, while interaction mainly affects the frequency, with some influence on the power as well. Enhancing the self-regulation weight of excitatory neurons and lowering that of inhibitory neurons significantly increases alpha power, revealing a potential mechanism for sex differences in alpha power. Additionally, enhancing the connection weight from inhibitory neurons to excitatory neurons, or decreasing the connection weight from excitatory neurons to inhibitory neurons, leads to an increase in both alpha peak frequency and alpha power.

Furthermore, we explored the impact of the E-I ratio on alpha rhythm characteristics (Figure 1F). In the neural circuit, the E-I ratio is defined in two ways: one for excitatory neurons, where it is calculated as W_EE_/W_EI_, and the other for inhibitory neurons, where it is calculated as W_IE_/W_II_. For both ratios, we simultaneously varied the respective variables (W_EE_ and W_EI_ for excitatory neurons; W_IE_ and W_II_ for inhibitory neurons), keeping the other two variables constant, and performed simulations. Figure 1GH shows how the alpha peak frequency and power change as the weights of these two connection types are altered. We found that for excitatory neurons, as the E-I ratio increased, the frequency decreased and the power increased. In contrast, for inhibitory neurons, as the E-I ratio increased, the frequency increased and the power decreased (Figure 1IJ).

In summary, for the E-I balance in neural circuits, inhibition leads to a decrease in power and an increase in frequency, while excitation only leads to an increase in power. The overall functioning of the neural circuit integrates the E-I balance of different neuron types. In certain neuronal systems, when excitation in excitatory neurons and inhibition in inhibitory neurons occur simultaneously, it is possible to observe an increase in both power and frequency. In actual experiments, under what conditions might such changes occur? One possibility is during the lifespan development process, as studies in different species have validated that from childhood to adulthood, both excitation and inhibition in the nervous system significantly increase, and then significantly decrease again from adulthood to old age. If this hypothesis holds true, it would also manifest in the frequency and power of alpha rhythms, exhibiting a pattern of first increasing and then decreasing. In the following results, we will use real EEG data to validate this hypothesis based on our computational simulation findings.

### Age dependent alpha-band activity in population level

To test the prediction from above simulation, we combined the three datasets (HBN, MPI and SRM) with a total of 3265 participants. These datasets share 58 common EEG electrode locations. We then obtained the spectra for each electrode in the eyes-closed (EC) state. Clear peaks were observed in both the alpha band across different age groups and genders. The dataset was separated with thirteen age groups (see methods), spectral profiles of males and females in different age ranges are clear and changing gradually (Figure 2A). As a typical example, using an electrode in the parietal region (P2), we narrowed the spectrum to the alpha frequency band to more clearly observe the characteristics of the alpha rhythm (Figure 2AB). At the population level, we found that both the alpha rhythm power and peak frequency exhibited age-dependent characteristics, regardless of sex (Figure 2C). In childhood, both alpha power and peak frequency were relatively low. These values peaked during puberty and early adulthood, and then declined in old age, forming an inverted U-shape. Surprisingly, the sex differences in alpha power and peak frequency were mainly observed during puberty and old age, with little difference during childhood and early adulthood, which showed a N-shape pattern (Figure 2D).

**Figure 2.**
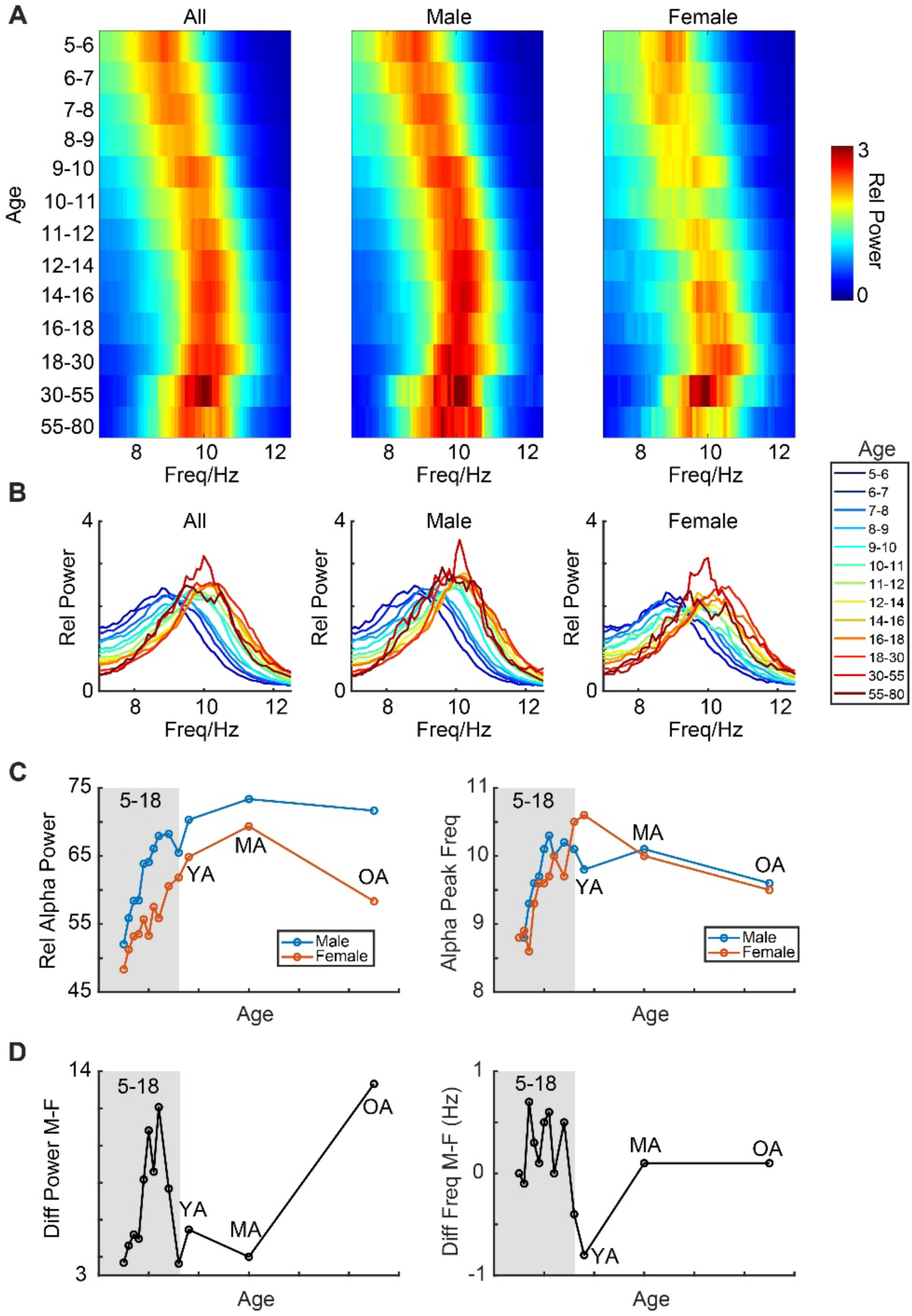
**Age and sex -dependent alpha rhythms in a typical electrode (P2)** A. Averaged relative power spectrum of all subjects (left), male (middle) and female (right) in closed (black curve) state in thirteen age groups in electrode P2. X axis is the frequency, y axis is the age group, color is the relative power. B. Same as A, with different type of demonstration. X axis is the frequency, y axis is the relative power, power spectrum with difference age groups is shown with different colors.C. Relationship between age and relative alpha power (left), and alpha peak frequency (right) in population level in electrode P2. D. Relationship between age and sex difference of relative alpha power (left), and sex difference of alpha peak frequency (right) in population level in electrode P2.

**Figure 3.**
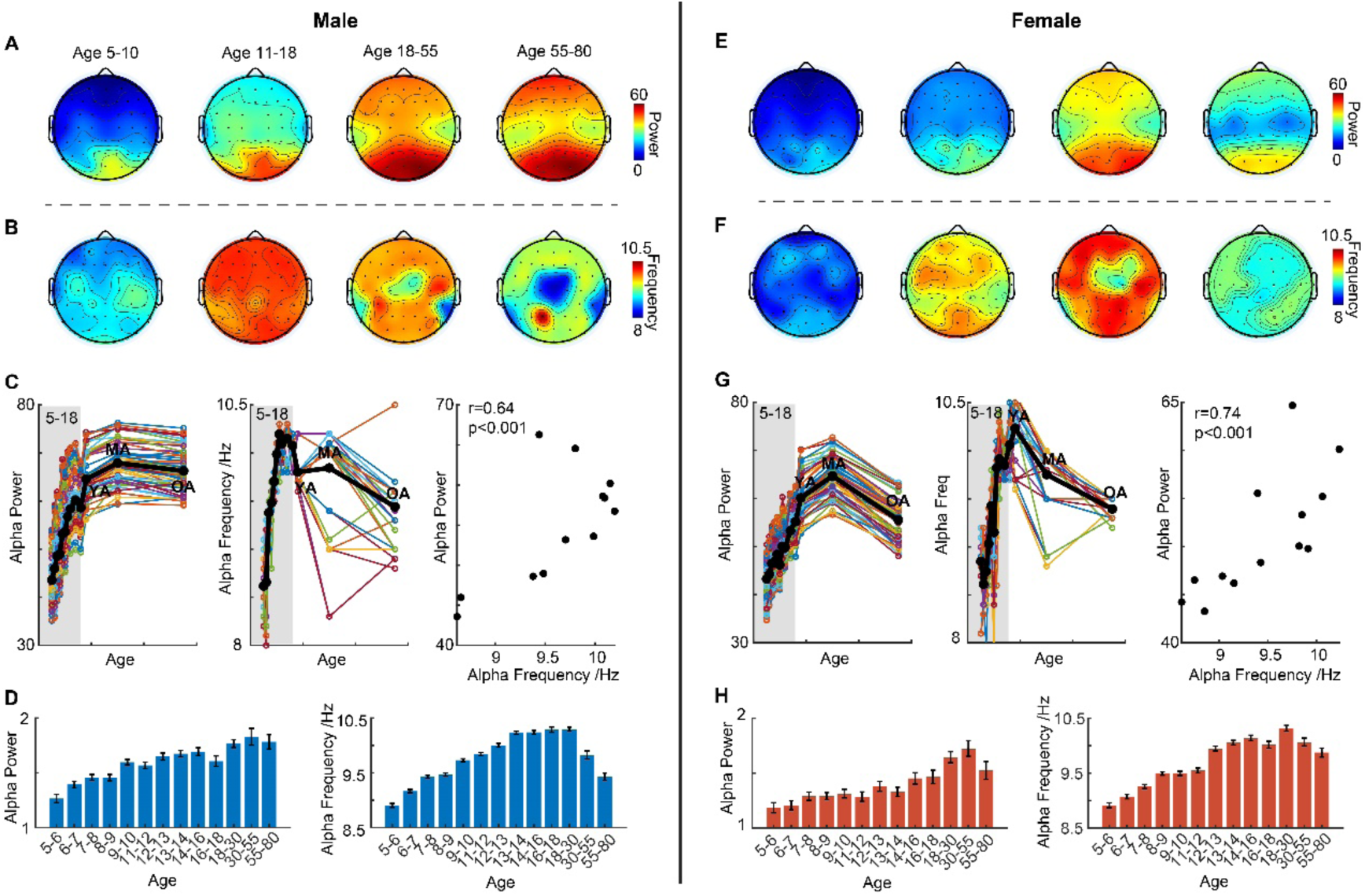
**Age-dependent features in alpha-band activity for males and females** A. Topographic maps of averaged alpha power for males in different age groups in eyes-closed states. B. Topographic maps of averaged alpha peak frequency for males in different age groups in eyes-closed states. C. Relationship between age and relative alpha power (left), and alpha peak frequency (middle) in population level for males in all electrodes. Averaged curve is shown by black curve. Relationship between alpha power and alpha peak frequency in population level is shown in right panel. D. Bar graphs with standard errors of alpha power (left) and peak frequency (right) for males in different age groups. E. Topographic maps of averaged alpha power for females in different age groups in eyes-closed states. F. Topographic maps of averaged alpha peak frequency for females in different age groups in eyes-closed states. G. Relationship between age and relative alpha power (left), and alpha peak frequency (middle) in population level for females in all electrodes. Averaged curve is shown by black curve. Relationship between alpha power and alpha peak frequency in population level is shown in right panel. H. Bar graphs with standard errors of alpha power (left) and peak frequency (right) for females in different age groups.

Further analysis of the alpha rhythm across all electrodes (Figure 3) revealed that alpha power was strongest in the occipital-parietal regions across all age groups (Figure 3AE). In contrast, alpha peak frequency did not have a specific region and its spatial distribution was not confined to the occipital-parietal region, but rather was closer to a whole-brain distribution (Figure 3BF). In Figure 3C and 3G, we show how alpha power (Figure 3CG left) and peak frequency (Figure 3CG middle) across all EEG electrodes change with age for both males and females (black line indicates the average). Both follow an inverted U-shape curve, with a significant positive correlation (Figure 3CG right). However, we also observed some differences. The alpha peak frequency starts to decline after reaching its peak in young adulthood (Figure 3DH left), which occurs earlier than the decline in alpha power, which peaks in middle adulthood and then decreases (Figure 3DH right). The different topographic maps and age curves suggest that changes in alpha power and alpha peak frequency are regulated by different neural mechanisms.

### Distinct time course of age dependent sex difference of alpha-band activity

Furthermore, we also compared the sex differences in alpha power (Figure 4A) and alpha peak frequency (Figure 4B) across different age groups. For alpha power, it is noteworthy that at age 5, there were no significant sex differences. These differences first appeared at age 6, initially in a small region of the occipital lobe. As age increased, the significant regions also expanded. Interestingly, from age 9 onwards, another significant sex difference gradually appeared in the bilateral parietal regions. These differences disappeared after age 16, and did not reappear until age 55, when differences emerged again in the bilateral parietal regions, while the occipital region showed no further differences. For alpha peak frequency, apart from around age 10 in the bilateral parietal regions, significant differences were not very strong in most age groups. Further, we fixed three brain region locations (Figure 4C) and conducted statistical analysis using two-way ANOVA for each region. The interaction effect was significant (p<0.001). We found that alpha power in the occipital and bilateral parietal regions represented two distinct components of sex differences. The occipital region reflected sex differences in early childhood (starting at age 6), while the parietal region reflected sex differences later in childhood (starting at age 9), with these differences reappearing after age 55. The timeline of this significant difference is more clearly depicted in Figure 4D, illustrating its different processes.

**Figure 4.**
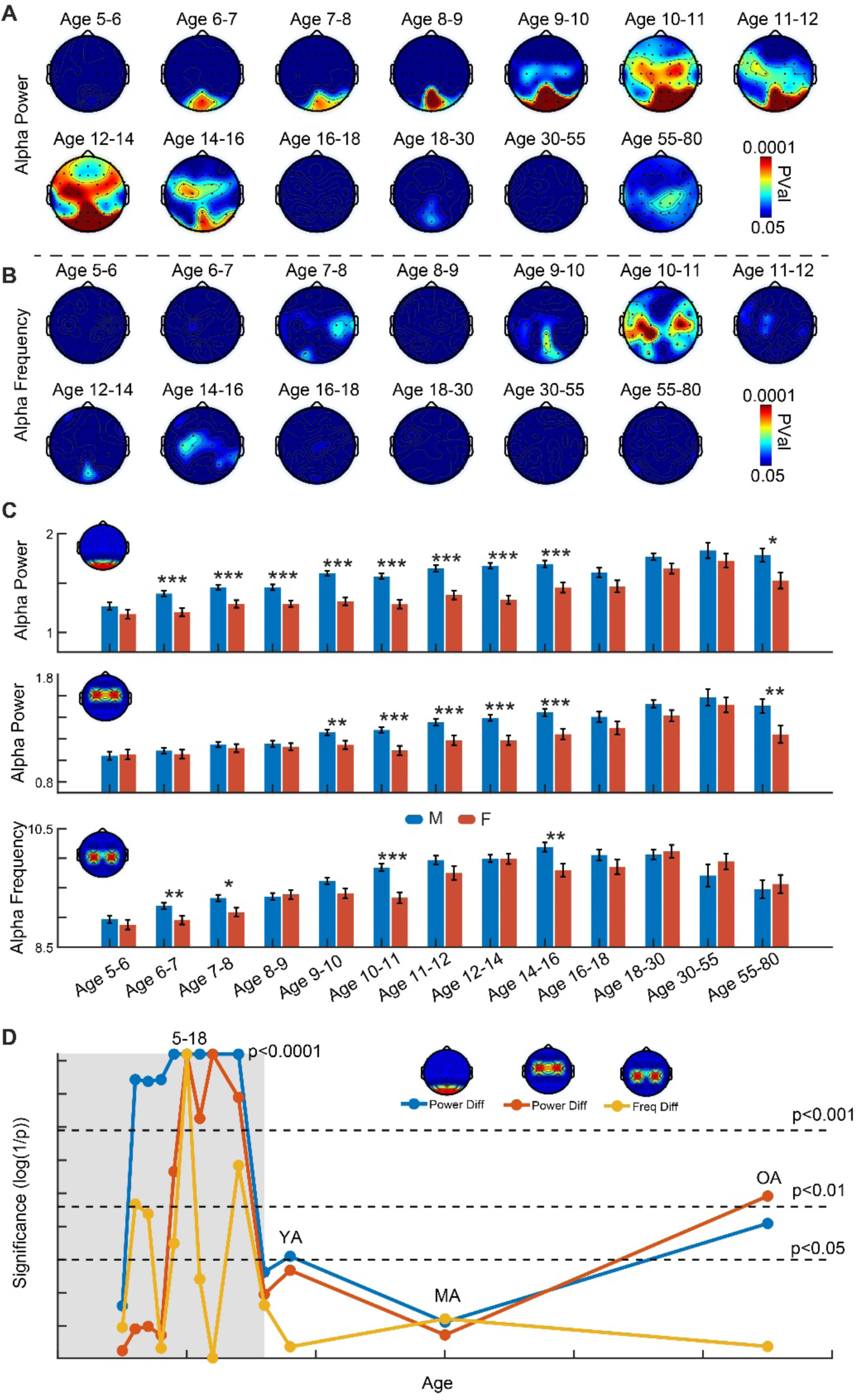
**Age dependent sex differences in different brain regions** A. Topographic maps of averaged alpha power for males in different age groups in eyes-closed states. B. Topographic maps of averaged alpha peak frequency for males in different age groups in eyes-closed states. C. Sex difference across different age groups in three typical brain regions. D. Different time courses of significance of sex difference across different age groups in three typical brain regions.

### Periodic activity contributes to age dependent sex difference of alpha-band activity

Next, we decomposed the alpha band into its periodic and aperiodic components to further explore their respective contributions. In Figure 5A, we present the decomposition results for different sex and age groups, where the pattern is clearly visible. For aperiodic activity, we extracted the power from the alpha band and found that, with increasing age, it did not follow an inverted U-shape curve. Instead, it exhibited an exponential decay pattern (Figure 5B Left), which is completely different from the phenomena observed in Figure 2. This also indicates that the results we previously found were not contributed by the non-periodic components in the spectrum. For periodic activity, it aligns completely with the earlier findings, whether in terms of alpha power (Figure 5B Middle) or alpha peak frequency (Figure 5B Right), both of which follow an inverted U-shape curve with age. The sex differences in alpha power and alpha peak frequency also show a similar N-shape curve (Figure 5C). This result further confirms that the periodic and non-periodic activities in the spectrum arise from different neural mechanisms.

**Figure 5.**
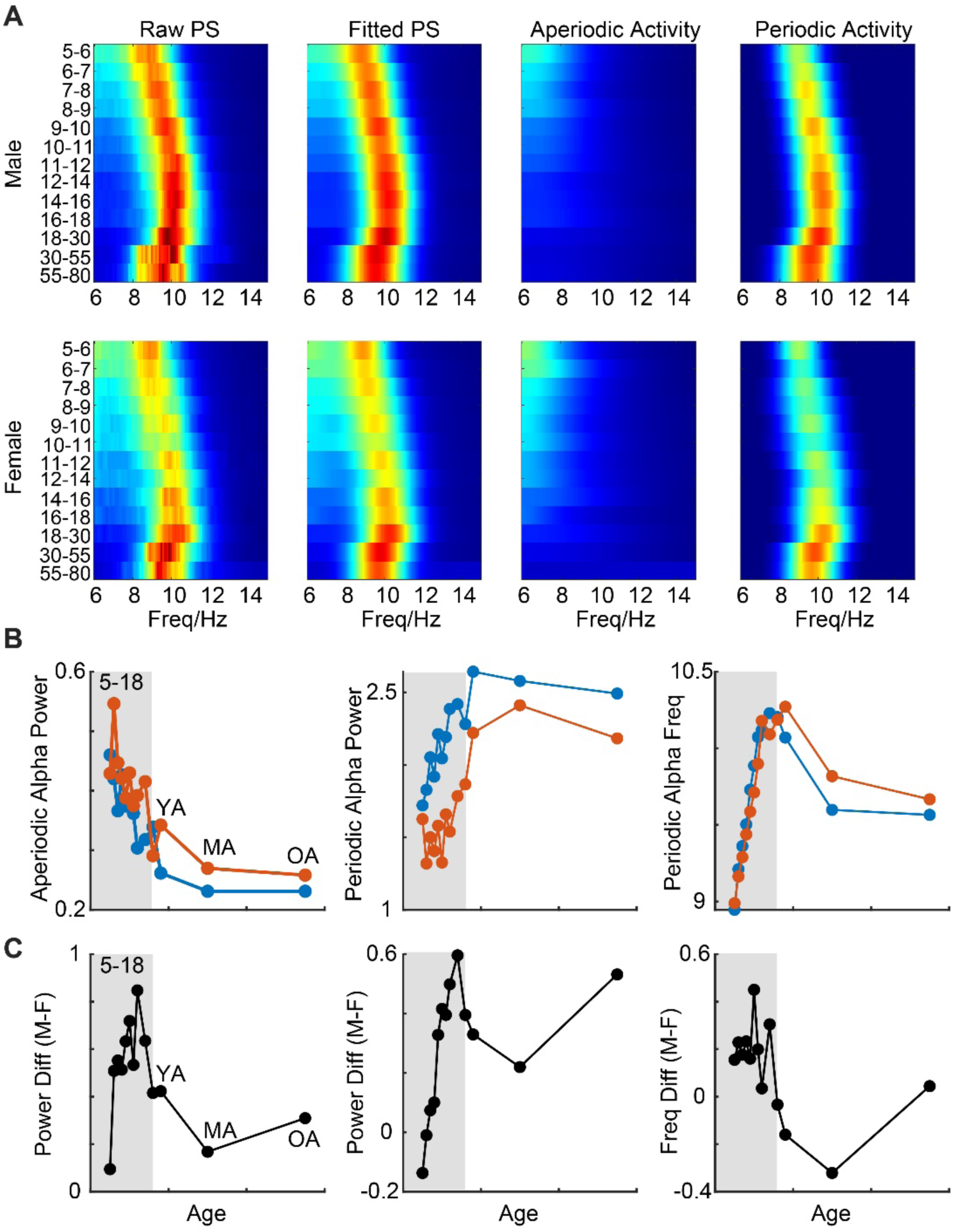
**Different contributions for aperiodic and periodic activity after dissecting** A. Dissecting periodic and aperiodic activity in population level in male and female. B. Relationship between age and aperiodic alpha power (left), periodic alpha power (middle) and periodic alpha peak frequency (right) in population level. C. Relationship between age and sex difference of periodic alpha power in occipital region (left), in parietal region (middle), and sex difference of periodic alpha peak frequency (right) in parietal region in population level.

## Discussion

In this study, we systematically depict for the first time the different trajectories of alpha-band activities (power and peak frequency) throughout the human lifespan in population level. We also compare the sex differences in these features at different age stages. By combining comparisons of brain topography, the decomposition of periodic and aperiodic components, and simulations of dynamic models, we reveal the intricate and distinct neural mechanisms through which age and sex influence the oscillatory characteristics of the alpha rhythm. The excitatory influence in self-regulation determines the sex differences in alpha power, while the inhibitory influence in interaction governs the age-related changes in both alpha power and alpha peak frequency. This significantly deepens our understanding of the alpha rhythm and lays a solid theoretical foundation for potential future applications.

### A framework of healthy oscillatory trajectory pattern in different age stages

Previous research in this field has primarily focused on specific age groups rather than the entire lifespan, such as studies on children to adolescents^86–90^ or young adults to older adults^91–95^. Our study effectively takes these fragmented pieces and puts them together to form a complete picture. One significant contribution of this research is that it clearly describes how the characteristics of alpha rhythm develop throughout a typical lifespan, from childhood to old age. The differences in alpha rhythm characteristics across age groups are substantial. Here, we emphasize that alpha power or alpha peak frequency should not be compared across groups without considering the age factor, as age is a key factor influencing these changes. When comparing alpha oscillation characteristics across different experimental groups, age must be considered, and efforts should be made to restrict the age range as much as possible. Additionally, specifically, the developmental trajectories of alpha power and alpha peak frequency differ, indicating that they are governed by distinct neural mechanisms. This was also confirmed in our model simulations (Figure 5), where alpha power is related to the excitatory self-regulation, and alpha peak frequency is linked to the inhibitory influence of interaction.

The study of these mechanisms has gained significance not only for advancing our knowledge of brain development and aging but also for potential clinical applications^96^ in understanding developmental mental and neurodegenerative disorders. When comparing healthy individuals with those suffering from psychiatric or neurological disorders, we may need sufficient alpha power to maintain brain health. In childhood, children with autism have significantly lower alpha power than normal controls^97–101^. In adulthood, individuals with ADHD also show significantly lower alpha power compared to normal controls^11,16^. In the elderly, both the AD and Stroke groups exhibit significantly reduced alpha power. Moreover, faster alpha peak frequency is generally associated with better brain health. In adolescence, the fetal alcohol spectrum disorder group shows a significant reduction in alpha peak frequency^90^. In adulthood, individuals with Schizophrenia also show significantly lower alpha peak frequency^96,102–105^. Similarly, in the elderly, the stroke^106–108^, Alzheimer’s diseases^22,109–111^ and Aneurysmal Subarachnoid Hemorrhage^81^ groups show marked decreases in alpha peak frequency. These may underscore the importance of understanding how the resting-state alpha rhythm evolves across the lifespan, as it could provide insights into the neurophysiological markers of brain function and dysfunction.

In addition, while much of the research on alpha rhythms has focused on the overall activity within the alpha frequency band, recent work suggests that alpha oscillations may consist of distinct periodic and non-periodic components^1,2^. These components could contribute differently to the observed patterns of alpha activity, making it important to decompose the alpha band and investigate their respective roles in age- and sex-dependent changes. Understanding the differential contributions of periodic and non-periodic components could offer valuable insights into the neural processes that underlie the modulation of alpha rhythms.

### Mechanisms of age-dependent sex difference in alpha-band activity

Another important contribution of this study to the field is its clear description of how the characteristics of alpha rhythm, namely sex differences, develop from childhood to old age. Both genders exhibit an inverted U-shape pattern over time, which further supports the idea that such trajectories are a universal pattern throughout the human lifespan. However, during the developmental process, we observed that the sex differences in the occipital and bilateral parietal regions follow different temporal patterns. This may be closely related to factors such as the differing rates of developmental maturation between genders and changes in hormones over time^112,113^. While some causal explanations cannot yet be fully established, the observed sex differences further highlight the need for a more nuanced understanding of how biological sex influences neural oscillatory patterns, especially during critical periods of development and aging.

The significant sex differences observed in both alpha power and peak frequency further complicate the understanding of these oscillatory patterns. Our analysis revealed that sex differences in alpha rhythms were most prominent during puberty and old age, with relatively small differences in childhood and young adulthood. This pattern aligns with the notion that hormonal changes during these life stages might influence the way alpha rhythms manifest. The re-emergence of sex differences in old age could reflect the interaction of hormonal changes with age-related neural degeneration, highlighting the dynamic nature of alpha oscillations over the lifespan. Moreover, the topographic analysis of alpha rhythms showed that the occipital and bilateral parietal regions exhibited distinct patterns of sex differences, further suggesting that different neural mechanisms govern alpha power and peak frequency in various brain regions. While occipital regions reflect early childhood sex differences (starting at age 6), the parietal regions are associated with later childhood differences (beginning at age 9) and show a re-emergence of sex differences after age 55. These regional differences indicate that the neural substrates underlying alpha rhythms are not uniform across the brain and are subject to developmental, aging, and gender-specific influences. In future experimental designs, researchers should also consider controlling for potential sex factors in specific age groups.

### Link between inhibition functions and alpha-band activity

The differential trajectories of these components suggest that they are governed by distinct neural mechanisms and contribute to different aspects of cognitive and developmental processes. A key observation is that alpha power and alpha peak frequency not only increase but also decrease during old age, indicating the reversibility of this process. In our simulation results (Figure 5), we found that the inhibitory function in the brain is likely key to altering these components. At younger ages, excitation and inhibition functions increase with age^114–128^; however, in older age, this inhibitory function becomes insufficient due to the loss of inhibitory neurons^129–132^, which has been widely validated across different species. Although there is currently no direct evidence linking behavior and alpha rhythm characteristics across the entire lifespan, this could serve as a direction for future research to further investigate.

Our results are also consistent with numerous previous studies on the functions of alpha rhythms^133–138^. In early childhood, alpha activity is often less pronounced, reflecting the developing brain’s ongoing organization and maturation. As individuals reach adolescence^139,140^, alpha rhythms become more prominent and are associated with improved attentional control, cognitive flexibility, and working memory. In adulthood^141–143^, enhanced alpha oscillations are often linked to a state of relaxed alertness and facilitating processes. With aging^109,144^, alpha activity tends to decrease in amplitude, which may reflect cognitive decline or reduced neural efficiency. The relationship between alpha oscillations and cognitive function thus appears to evolve with age, suggesting that changes in alpha rhythm are not only a marker of brain maturation but also a potential indicator of age-related cognitive changes. Understanding this dynamic relationship is crucial for exploring how neural oscillatory patterns contribute to cognitive abilities across the lifespan.

### Future Directions

Future studies would aim to explore the underlying neural mechanisms that drive the changes of alpha rhythm, including the role of neurotransmitters, hormonal influences, and structural changes in the brain. Investigating how these factors interact with alpha rhythms could provide valuable insights into the biological underpinnings of cognitive aging and neurodevelopmental processes. Additionally, longitudinal studies examining these dynamics across the lifespan will be crucial for further clarifying the role of alpha oscillations in human brain development and aging.

## Acknowledgments

This work was funded by the Research Grants Council of the Hong Kong Special Administrative Region, China (Project Nos. R4022–18, N_CUHK456/21, 14114721, 14119022, and STG1_M-401_24-N to V.C.K.C. Project No. 9042986 to R.H.M.C.), grants from The Chinese University of Hong Kong (Project Nos. 2020095, 2021.065 for Faculty of Medicine, and project “Impact Case C7” for Research Committee, to V.C.K.C.), and funding from City University of Hong Kong (No. 7005641, 7005857 and 7006091 to R.H.M.C.)

## Conflicts of interest statement

The co-authors declare that the research was conducted in the absence of any commercial or financial relationships that could be construed as a potential conflict of interest.

## Contributors

CH, RC and VC conceived and designed the study. CH contributed to the literature search, data analysis, and the interpretation of results. All authors contributed to writing the paper.

## Notes

### Competing Interest Statement

The authors have declared no competing interest.

